# SARS-CoV-2 triggered excessive inflammation and abnormal energy metabolism in gut microbiota

**DOI:** 10.1101/2021.11.08.467715

**Authors:** Tuoyu Zhou, Yufei Zeng, Jingyuan Wu, Junfeng Li, Jun Yan, Wenbo Meng, Hawen Han, Fengya Feng, Jufang He, Shuai Zhao, Ping Zhou, Ying Wu, Yanling Yang, Rong Han, Weiling Jin, Xun Li, Yunfeng Yang, Xiangkai Li

## Abstract

Specific roles of gut microbes in COVID-19 progression are critical. However, the circumstantial mechanism remains elusive. In this study, shotgun metagenomic or metatranscriptomic sequencing were performed on fecal samples collected from 13 COVID-19 patients and controls. We analyzed the structure of gut microbiota, identified the characteristic bacteria and selected biomarkers. Further, GO, KEGG and eggNOG annotation were employed to correlate the taxon alteration and corresponding functions. The gut microbiota of COVID-19 patients was characterized by the enrichment of opportunistic pathogens and depletion of commensals. The abundance of *Bacteroides* spp. displayed an inverse relationship to COVID-19 severity, whereas *Actinomyces oris*, *Escherichia coli*, and *Gemmiger formicilis* were positively correlated with disease severity. The genes encoding oxidoreductase were significantly enriched in SARS-CoV-2 infection. KEGG annotation indicated that the expression of ABC transporter was up regulated, while the synthesis pathway of butyrate was aberrantly reduced. Furthermore, increased metabolism of lipopolysaccharide, polyketide sugar, sphingolipids and neutral amino acids was found. These results suggested the gut microbiome of COVID-19 patients was correlated with disease severity and in a state of excessive inflammatory response. Healthy gut microbiota may enhance antiviral defenses via butyrate metabolism, whereas the accumulation of opportunistic and inflammatory bacteria may exacerbate the disease progression.

## 1. INTRODUCTION

The COVID-19 pandemic caused by SARS-CoV-2 triggered acute and severe respiratory pathology, but growing evidence suggested that complicating gastrointestinal symptoms is common as extrapulmonary manifestations [1]. SARS-CoV-2 was detected in both fecal and anal swab of infected persons [2], while high anal swab viral load had been associated with adverse clinical outcomes in patients. Further, some cases suggested that untreated sewage might increase the fecal-oral transmission risk of the virus [3, 4]. SARS-CoV-2 infects host cells through the ACE2 receptor [5], and continuously replicates in the gastrointestinal system [6], thereby weakening the intestinal barrier. It had been authenticated that ACE2 was a vital regulator of intestinal inflammation [7]. The deficiency of ACE2 may aggravate the gut microbiota imbalance and gastroenteritis-like symptoms by altering the inflammatory sensitivity [8].

Gut microflora provides various biological functions for the host, including promoting immune system homeostasis, metabolizing nutrients, and maintaining the intestinal mucosal barrier [9]. At the same time, microbial flora is also thought to be a contributing factor in virus clearance [10]. In contrast, the intestinal flora dysbiosis reduces antiviral immune responses and aggravated respiratory diseases [11]. Consumption of antibiotic-sensitive gut microbes could augment the susceptibility to pulmonary allergic inflammation and influenza virus infection [12, 13]. Severe influenza A virus infection was associated with intestinal disease and altered intestinal flora [14]. The greater abundance of *Escherichia coli* and *Enterococcus faecium* in the H7N9 patients might be account for bacteremia and abdominal infection [15].

Existing studies described the close link between microbial florae dysbiosis and SARS-CoV-2 infection [16]. Compared with healthy controls, COVID-19 patients showed significantly lower bacterial diversity [17], while opportunistic pathogens enrichment and beneficial bacteria depletion were also observed [17–19]. The consumptive beneficial commensals belong to the *Ruminococcaceae* and *Lachnospiraceae* [17, 18] , and the butyrate-producing bacterium *Facealibacterium prausnitizii* was found to be negatively associated with COVDI-19 severity [19, 20]. In contrast, the putrefactive bacterium *Coprobacillus* were correlated to the disease severity [18]. Notably, some specific *Bacteroides* spp., capable of down-regulating ACE2 expression in the murine gut, are inversely correlated with the SARS-CoV-2 load [18]. These results highlight the potential role of gut microbiota in the disease predisposition of COVID-19 patients. Nevertheless, the specific mechanism of interaction between SARS-CoV-2 and gut microbiota remains elusive. Especially, the association between taxon and related functions should be explored in depth.

The latest research suggested there may exist energy competition between SARS-CoV-2 infection and gut microbiota [21], while that microbiome dysbiosis and malnutrition were proved to be linked with COVID-19 susceptibility [19]. Therefore, we hypothesized that SARS-CoV-2 infection would alter the composition and function of gut microbiota, while the healthy intestinal flora may critically maintain immune homeostasis and energy supply against COVID-19 development [22]. Through metagenome (MG) and metatranscriptome (MT) sequencing, this study annotated the gut microbiome information of COVID-19 patients and described the alterations in core microbial communities with their related functions. Additionally, we revealed the connection between these changes and clinic features of COVID-19 patients, further elucidating their interactions in active transcripts features.

## 2. RESULT

### 2.1 Information of subjects

The age of 13 COVID-19 patients ranged from 21 to 50 years, with a median age of 24 years old. Most COVID-19 cases are male, accounting for about 76% (Table S1). 8 COVID-19 cases developed clinical symptoms such as fever or sore throat (Table S1), and 5 cases are asymptomatic carriers. The lymphocyte and APTT levels of COVID-19 patients were significantly lower than those in health controls, while hemoglobin and total bilirubin levels were significantly higher than those in health group (Fig. 1). Although most physiological indicators in CAP were similar with COVID-19 group, the globulin, D-Dimer and fibrinogen levels were significantly different (Fig. 1).

**Figure 1.**
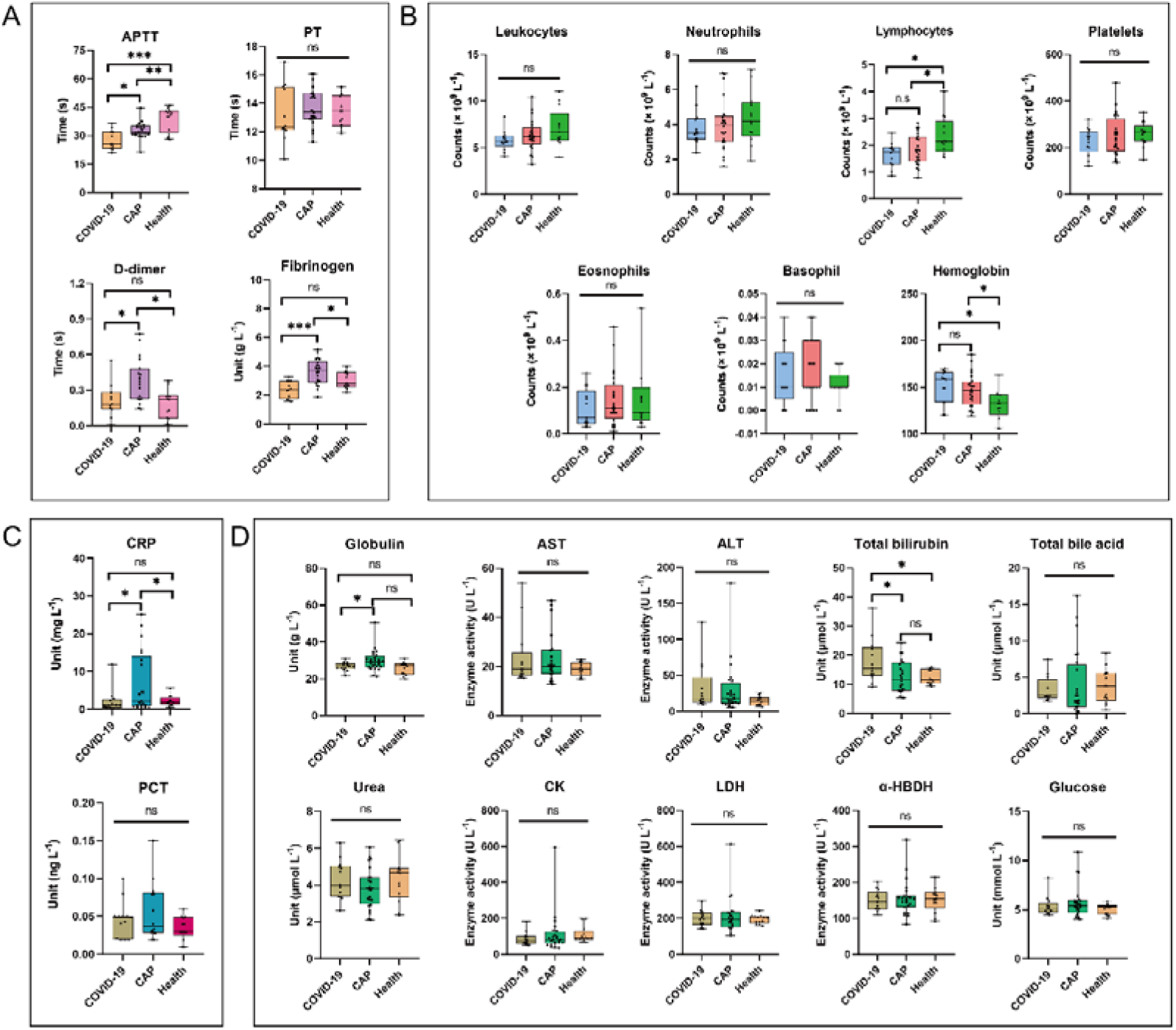
Comparation of clinical biomarkers between COVID-19 (n=13), heath (n=13) and CAP (n=25) groups, including (A) coagulation parameters; (B) blood routine results; (C) infectious criterions and (D) biochemistry indexes. PT, prothrombin time; APTT, activated partial thromboplastin time; CRP, c reactive protein; PCT, procalcitonin. AST, aspartate aminotransferase; ALT, alanine aminotransferase; CK, creatine kinase; LDH, lactate dehydrogenase; α-HBDH, α-hydroxybutyric dehydrogenase.

### 2.2 Gut microbiota structure dissimilarity among COVID-19, health, and CAP groups

9.31 to 11.79 G high-quality sequence reads were obtained from fecal samples of these three groups, using MG sequencing (Table S2). SARS-CoV-2 infection had the greatest impact on fecal mycobiome (PERMANOVA test, R^2^=0.06, P =0.02) of COVID-19 patients, while age, sex, antibiotics, and antiviral drugs had no significant impact (Fig. S3). Albeit the Shannon index and Chao index of the baseline samples in the COVID-19 group were close to those in the health group, these indexes based on the last follow-up samples were significantly decreased (Fig. 2A and Fig. 2B). PCoA plot indicated the fecal microbiome of healthy subjects clumped together, while the samples of COVID-19 group developed stronger heterogeneity (Fig. 2C and Fig. S4). Microbes present in all samples from each subject group were defined as core microbes. All three groups shared 1201 core microbes, while COVID-19 and the health groups owned 2241 common microbes (Fig. 2D). The individual difference in fecal bacterial community of COVID-19 group was about twice that in healthy controls (P =0.00036, Fig. S5A).

**Figure 2.**
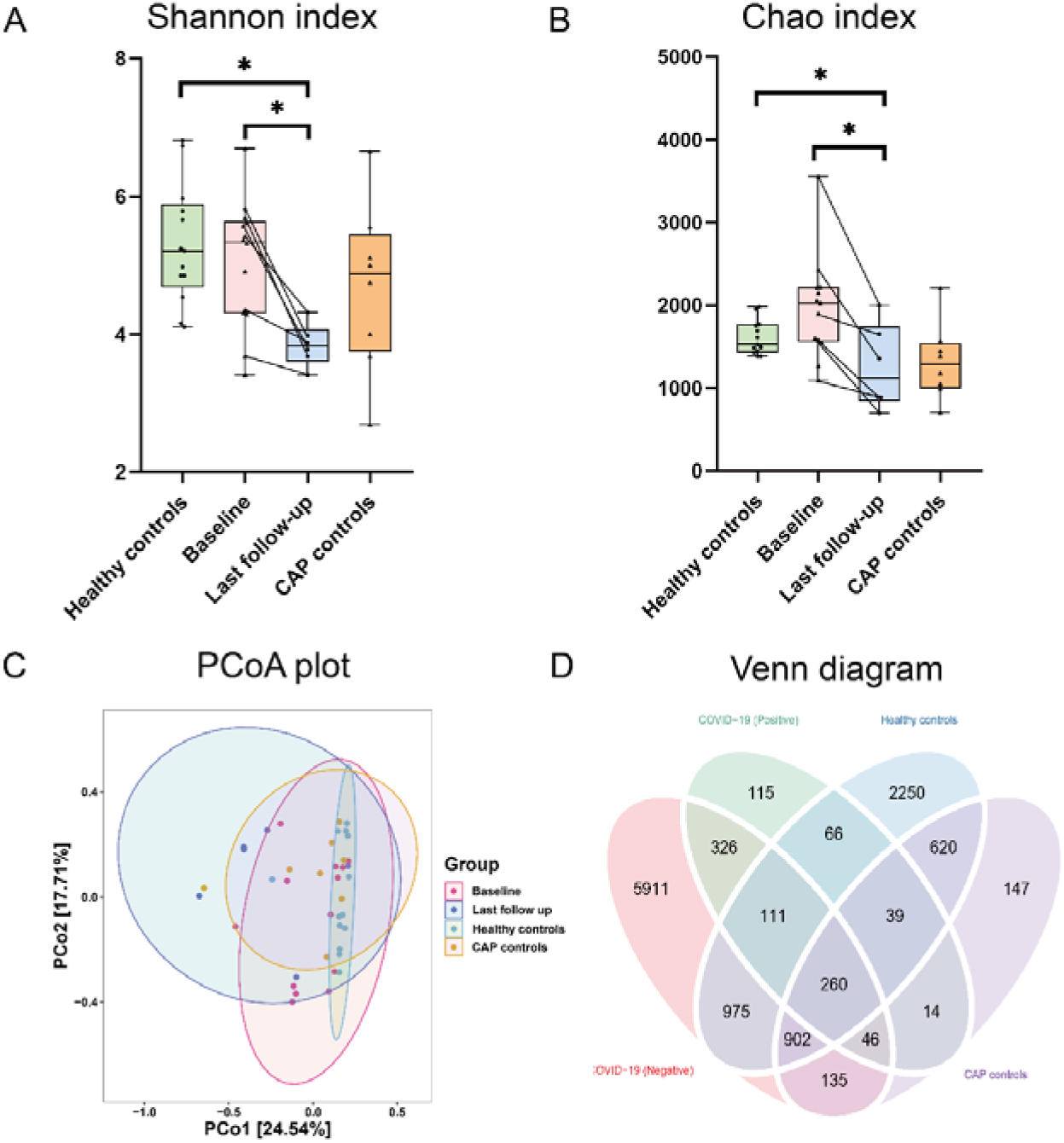
Alteration in gut microbial diversity and community structures in COVID-19 (n=13), healthy (n=13) and CAP (n=8) subjects. Alpha diversity of the gut microbiota among the three groups based on the (A) Shannon index and (B) chao index. (C) Microbiome communities were assessed by principal coordinate analysis (PCoA) of Bray-Curtis distances. (D) Venn diagram presenting the overlap of OTUs of the fecal microbiome across the three groups.

Subsequently, the effect of temporal succession on the COVID-19 microbiota was investigated. Baseline samples of 6 moderate/severe COVID-19 patients were inconsistent with those from healthy individuals, while the baseline samples from mild patients (except for COV11) were within normal intestinal microbiota variation (Fig. S5B). The bacterial communities of two COVID-19 patients (COV1, COV5) gradually recover to healthy level over time. Although SARS-CoV-2 was negative on throat swabs, the microbiomes of COV2, COV9, COV10, and COV11 was increasingly different from healthy controls (Fig. S5B).

### 2.3 Taxa composition of COVID-19, health, and CAP groups

To investigate the alteration of microbiota taxa, the relative proportion of microorganisms was assessed at the phylum, family, genus, and species levels (Fig. 3 and Table S4). Versus healthy ones, the relative abundance of phylum *Bacteroidetes* decreased in COVID-19 group, while *Actinobacteria* and *Verrucomicrobia* significantly enriched (Fig. 3A and Fig. S6A). The *Firmicutes*/*Bacteroidetes* (F/B) ratio of COVID-19 was higher than observed in health (P=0.04) and CAP groups (P=0.05) (Fig. S6B). The relative abundance of *Bacteroidaceae*, *Lachnospiraceae* and *Rikenellaceae* of COVID-19 group were lower than CAP and health groups, while *Actinomycetaceae* was greater than these two control groups (Fig. 3B). At the genus level, *Bacteroides, Faecalibacterium* and *Blautia* significantly decreased in COVID-19 group, but the *Escherichia, Gemmiger*, and *Subdoligranulum* were specially up-regulated. Intriguingly, *Akkermansia* expanded in both COVID-19 and CAP groups (Fig. 3C). Compared to healthy group, *Prevotella copri*, *Clostridium leptum*, *Alistipes putredinis* and *Prevotella* sp. BV3P1 consumed in COVID-19 group, while the relative abundance of *Gemmiger formicilis* remarkably increased. (Fig. 3D). The LEfSe results identified that *Rothia*, *Lactobaccillus*, *Schaalia*, *Actinomyces* and *Akkkermansia* dominated in the COVID-19 group (Fig. 4A, Fig. S7A and Table S5). The phylogenetic distribution profiles of COVID-19 group, and healthy group is non-overlapping (Fig. 4B). Spearman correlation analysis revealed a significant negative correlation between the characteristic bacteria in COVID-19 and health groups (Fig. 4C and Fig. S7B). The 20 most dominant species based on the random forest model (Fig. S8) were compared with the LEfse results, and two biomarkers (pathogens *Barnesiella* and *Chlamydia*) were obtained to distinguish COVID-19 group from health group (Fig. 4D). Besides, COVID-19 patients at the last follow up were further enriched with *Klebsiella pneumoniae*, *Escherichia coli*, *Shigella dysenteriae* and *Shigella flexneri* (Table S5). To assess the correlation between baseline fecal microbiota and COVID-19 severity, COVID-19 group was divided into mild and moderate/severe, using health group as baseline. Bacteria responsible for COVD-19 severity included *Escherichia coli*, *Burkholderiales* bacterium, *Actinomyces oris*, *Streptococcus parasanguini*, *Gemmiger formicilis* and *Eisenbergiella Tayi*. All the bacteria negatively related to COVID-19 severity originated from *Bacteroides* (e.g., *Bacteroides thetaiotaomicron*, *Bacteroides caccae* and *Bacteroides Fragilis* (Table 1).

**Figure 3.**
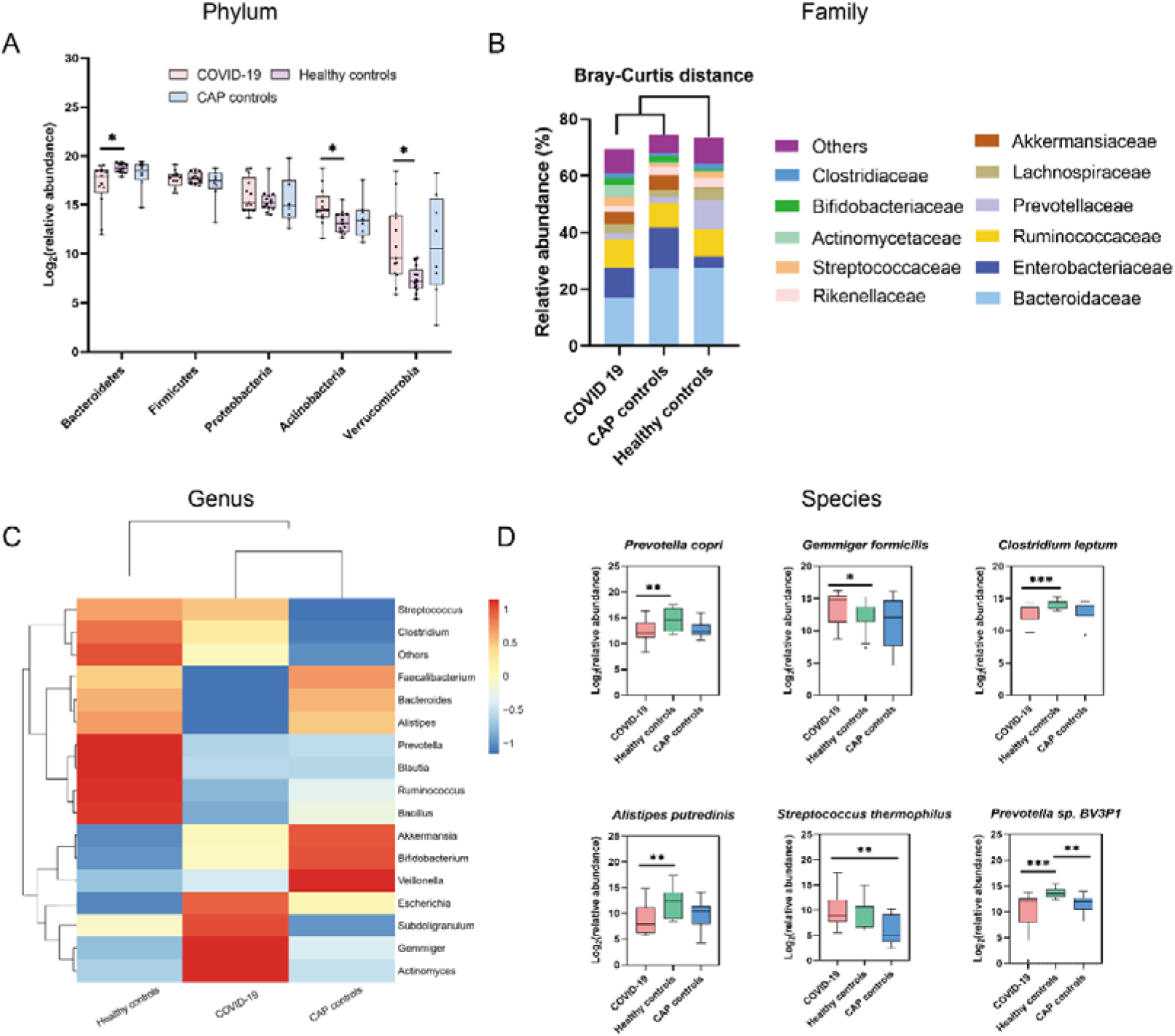
Taxonomic differences in the stool microbiota between COVID-19 patients, CAP patients, and healthy controls. Comparison of the relative abundance at the phylum (A), family (B), genus (C) and species (D) levels across the three groups. Specific to box figure, each box corresponds an interquartile range of taxa abundance, and the black line represents to median abundance. Vertical lines indicate the variability in the abundance of each taxon.

**Figure 4.**
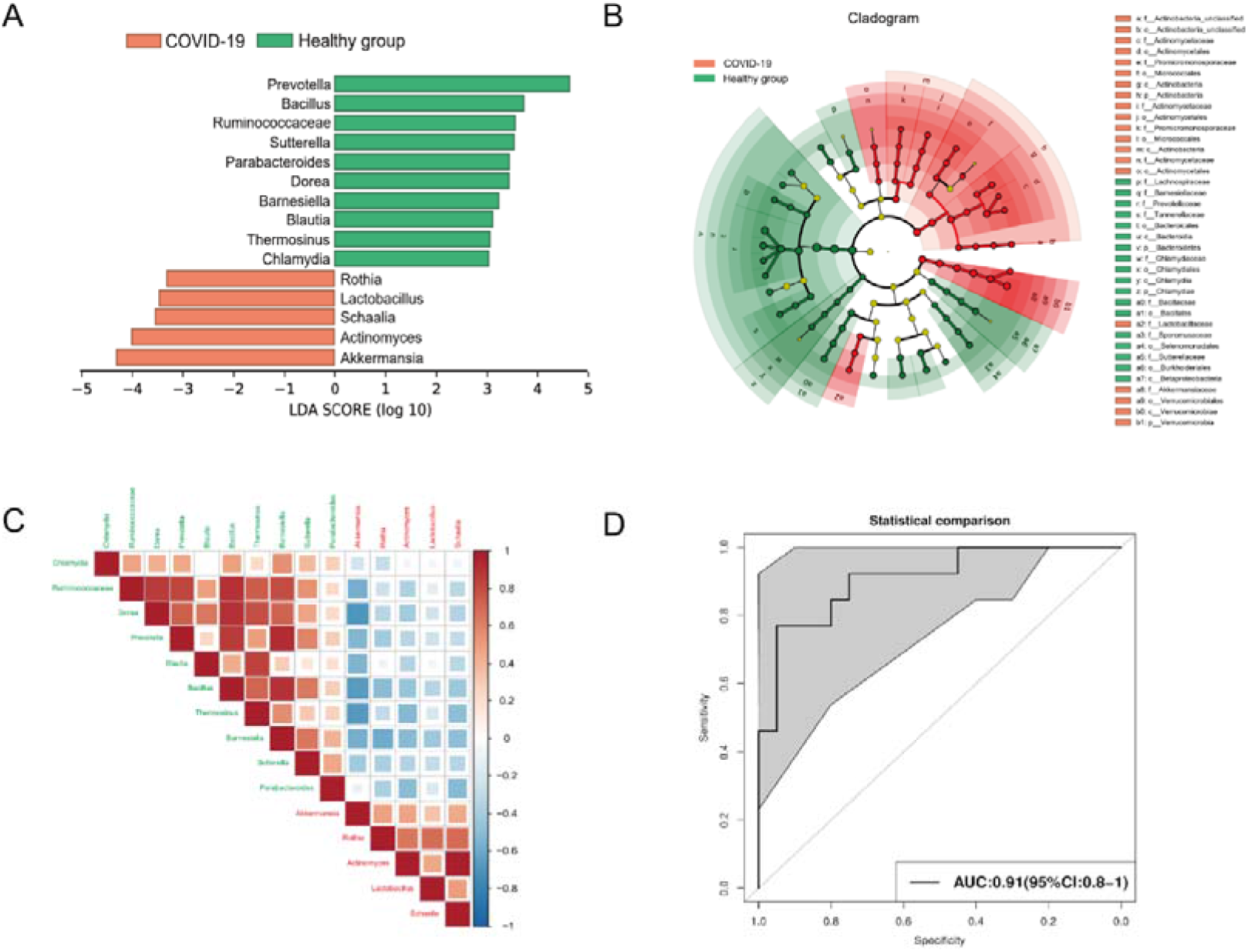
Taxonomic differences of fecal microbiome in COVID-19 and health group. (A) LEfSe analysis conducted to reveal the significant differences in microbiota composition between COVID-19 patients (orange) and healthy controls (emerald). (B) Cladogram using LEfSe method indicated the phylogenetic distribution of fecal microbiota associated with COVID-19 group and healthy subjects. (C) Spearman correlation of associated genera in COVID-19 patients and healthy controls. (D) Prediction of two biomarkers (*Barnesiella* and *Chlamydia*) in the microbiome of COVID-19 and healthy controls. The area under the ROC curve is displayed in the center. p, phylum; c, class; o, order; f. family; g, genus.

**Table 1.**
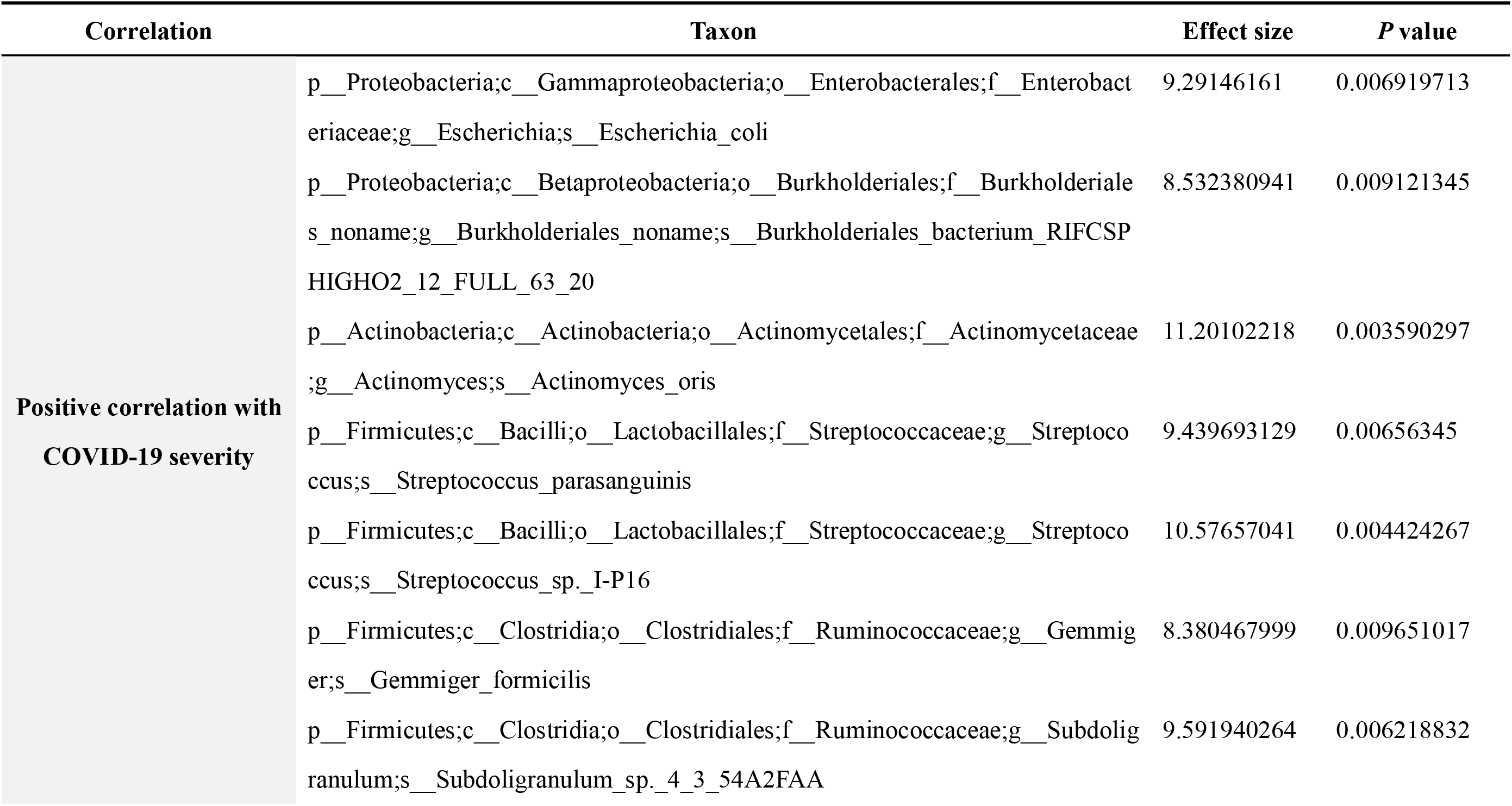
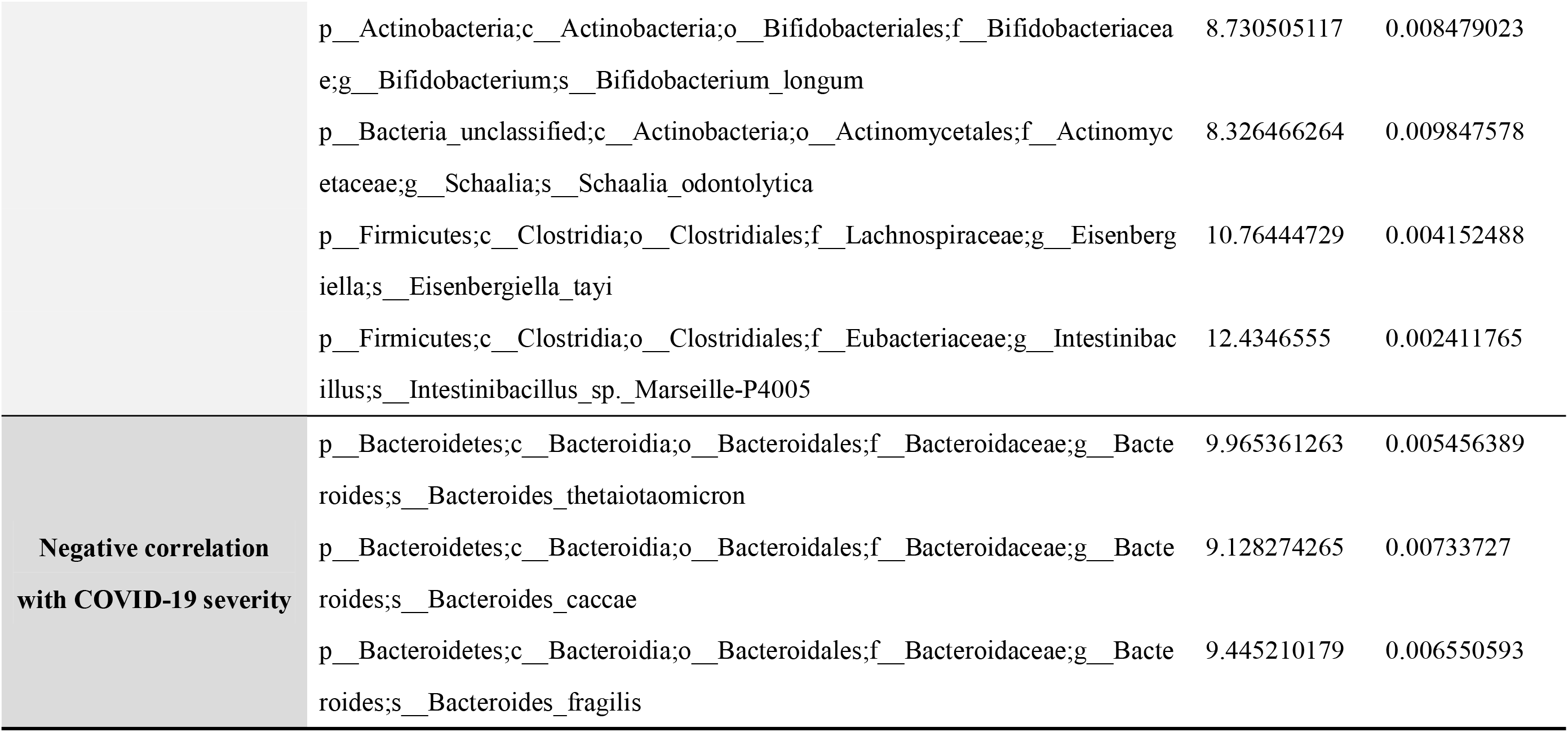
Intestinal bacteria associated with COVID-19 severity.

### 2.4 Functional characteristics of gut microbiome in COVID-19 group

Compared with health group, GO classification demonstrated the genes with RNA-mediated transposition, growth, and transport were significantly up-regulated in the molecular function category. Genes related to the integral component of membrane and cell wall were especially enriched in cellular component category. According to the biological process category, genes involved in the protein binding, single-stranded RNA binding, and structural constituent of ribosome were significantly increased (Fig. 5A). SARS-CoV-2 infection also caused remarkably alteration in metabolic pathways (Fig. 5B). Most of them were relevant metabolism processing (namely, Tryptophan metabolism; Sphingolipid metabolism; Galactose metabolism, etc.), followed by Human disease (*Staphylococcus aureus* infection; *Salmonella* infection, etc.) Bacterial invasion of epithelial cells), Environmental Information Processing (Bacterial secretion system, ABC transporters), Genetic Information Processing (CAMP resistance and β–Lactam resistance) and Cellular Processes (Biofilm formation – *Escherichia coli*). Notably, significant change was found in energy metabolism of COVID-19 group ((Fig. S10), and pathway entry also indicated that the butyrate synthesis pathway was remarkably lower than that in health group (Fig. S11). Furthermore, compared with CAP group, KEGG pathways enriched in the COVID-19 group were involved in opportunistic pathogen *Pseudomonas aeruginosa* and *Staphylococcus aureus* infection (Fig. S9).

**Figure 5.**
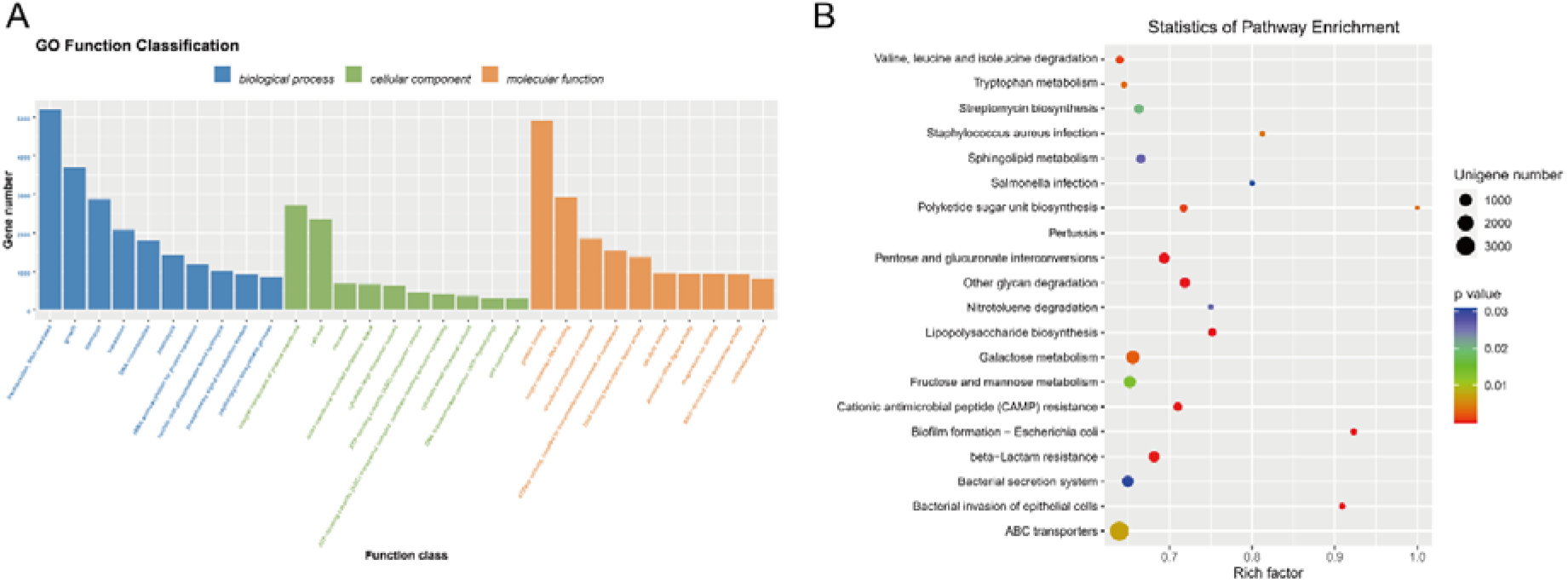
(A) Functional classification of differential genes up-regulated in the COVID-19 group according to Gene ontology (GO) terms in the domains ‘molecular function’ (MF) ‘cellular component’ (CC) and ‘biological process’ (BP). (B) Statistics of Kyoto Encyclopedia of Genes (KEGG) annotation of differential genes up-regulated in the COVID-19 group. The size of each circle represents the number of significant unigenes up regulated in the corresponding pathway (the significant threshold of differential genes as an absolute value of log2 (fold change) ≥1, p <0.05). The up-regulation factor was calculated with the number of up-regulated gene divided by the total number of background genes in the corresponding pathway. A pathway with a p-value <0.03 is considered significantly over-represented.

### 2.5 Comparation of taxonomic and functional differences between MG and MT in COVID-19 group

To examine the potential activity of intestinal microbes detected in COVID-19 patients , 10 COVID-19 baseline fecal samples underwent MT sequencing and an average of 16.78 G reads was generated (Table S3). The MG data showed lower alpha diversity than that in MT data (Fig. S12). Five major phyla identified in MG were also confirmed in MT (Fig. 6A). *Bacteroides* and *Alistipes* were the most main bacterial genera, while *Clostridium leptum* and *Bacteroides vulgatus* were the most abundant species in MG (Fig. 6B and 6C). In MT, *Bacteroides* and *Prevotella* were main bacterial genera, with highest abundance of *Gemmiger formicilis* and *Faecalibacterium prausnitzii* (Table S6 and S7). Next, we analyzed the ratio of the mean relative abundance in the MG to those in the corresponding MG (MT /MG ratio) to explore the relative activity of the baseline COVID-19 microbiome (Fig. 6D). The results demonstrated the relative activity of some beneficial bacteria, including *Blautia*, *Alistipes*, *Clostridium leptum* and *Akkermansia muciniphila* [23], was decreased, while some pernicious bacteria, such as *Prevotella copri* [24], *Subdoligranulum*, *Escherichia coli* and *Gemmiger formicilis* displayed high transcriptional activity. It is noteworthy that, a high MT/MG ratio of several bacteria negatively correlated with COVID-19 (i.g., *Faecalibacterium prausnitzii, Bacteroides ovatus*, *Bacteroides fragilis* and *Bacteroides caccae*) were observed (Table S8).

**Figure 6.**
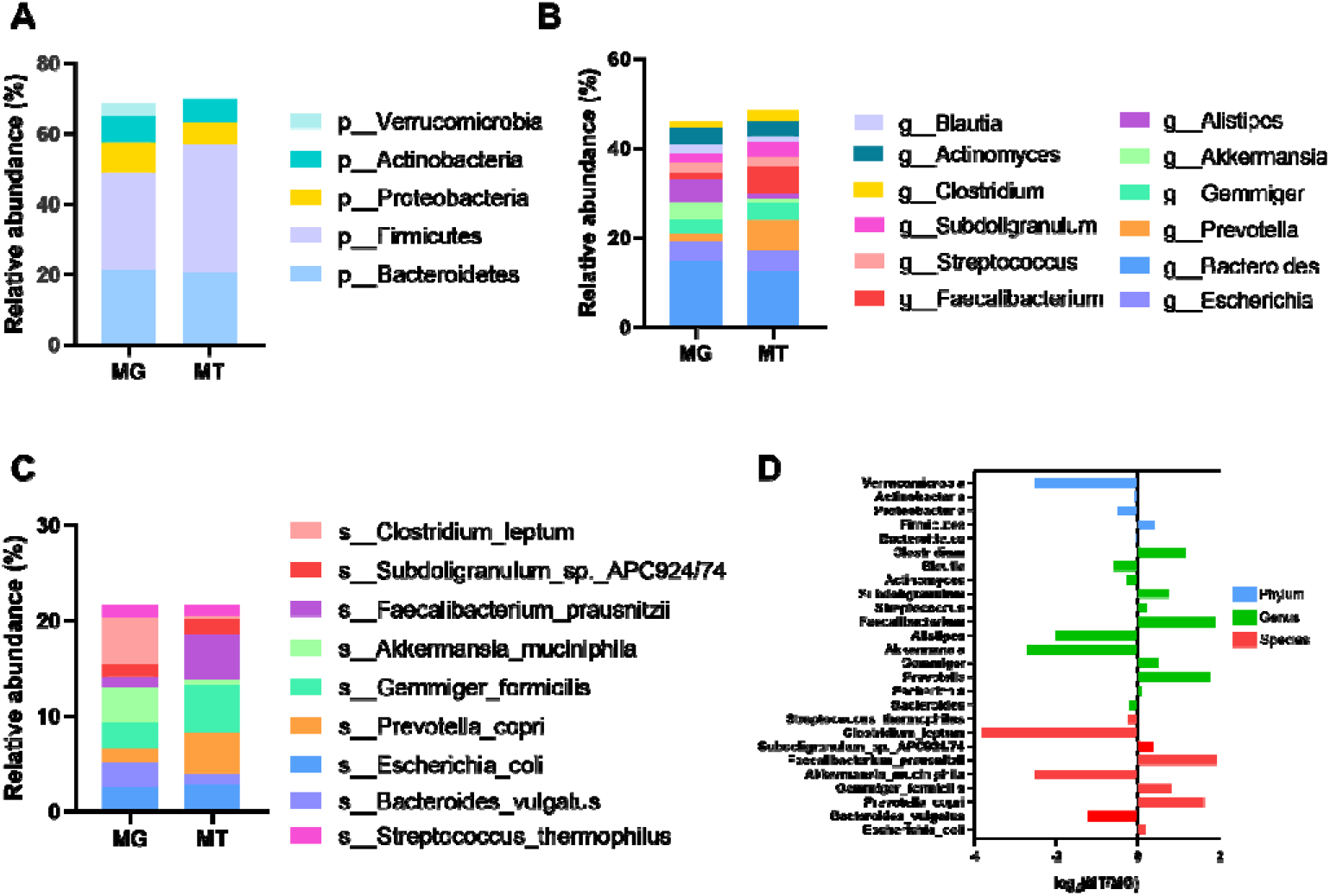
Gut microbiome composition of metagenome (MG) and metatranscriptome (MT) in COVID-19 patients. (A–C) illustrates bacterial composition at phylum, genus and species levels (relative abundance ≥1%), respectively. Microbiota composition indicates the average relative abundance of bacterial presence in all samples (n=10). (D) Ratio of mean relative abundance of microbes in MT to that in MG (MT/MG).

To functionally characterize the active gut microbiome of COVID-19 patients, unigenes of the MG and MT were aligned to protein sequences from KEGG and eggNOG databases. Of the MG data, KEGG pathways pertaining to ‘Carbohydrate metabolism’, ‘Amino acid metabolism’ and ‘Membrane transport’ were noticeably up regulated (Fig. 7A). Likewise, microbial genes involved in ‘Amino acid transport and metabolism’, ‘Replication, recombination and repair’ and ‘Carbohydrate transport and metabolism’ pathways were highly enriched in EggNOG functional categories (Fig. 7C). For MT data, the highly enriched KEGG pathways involved in ‘Carbohydrate metabolism’, ‘Amino acid metabolism’, ‘Nucleotide metabolism’ and ‘Membrane transport’ (Fig. 7B). The most active eggNOG functional categories were ‘Replication, recombination and repair’ ‘Carbohydrate transport and metabolism’, followed by ‘Amino acid transport and metabolism (Fig. 7D).

**Figure 7.**
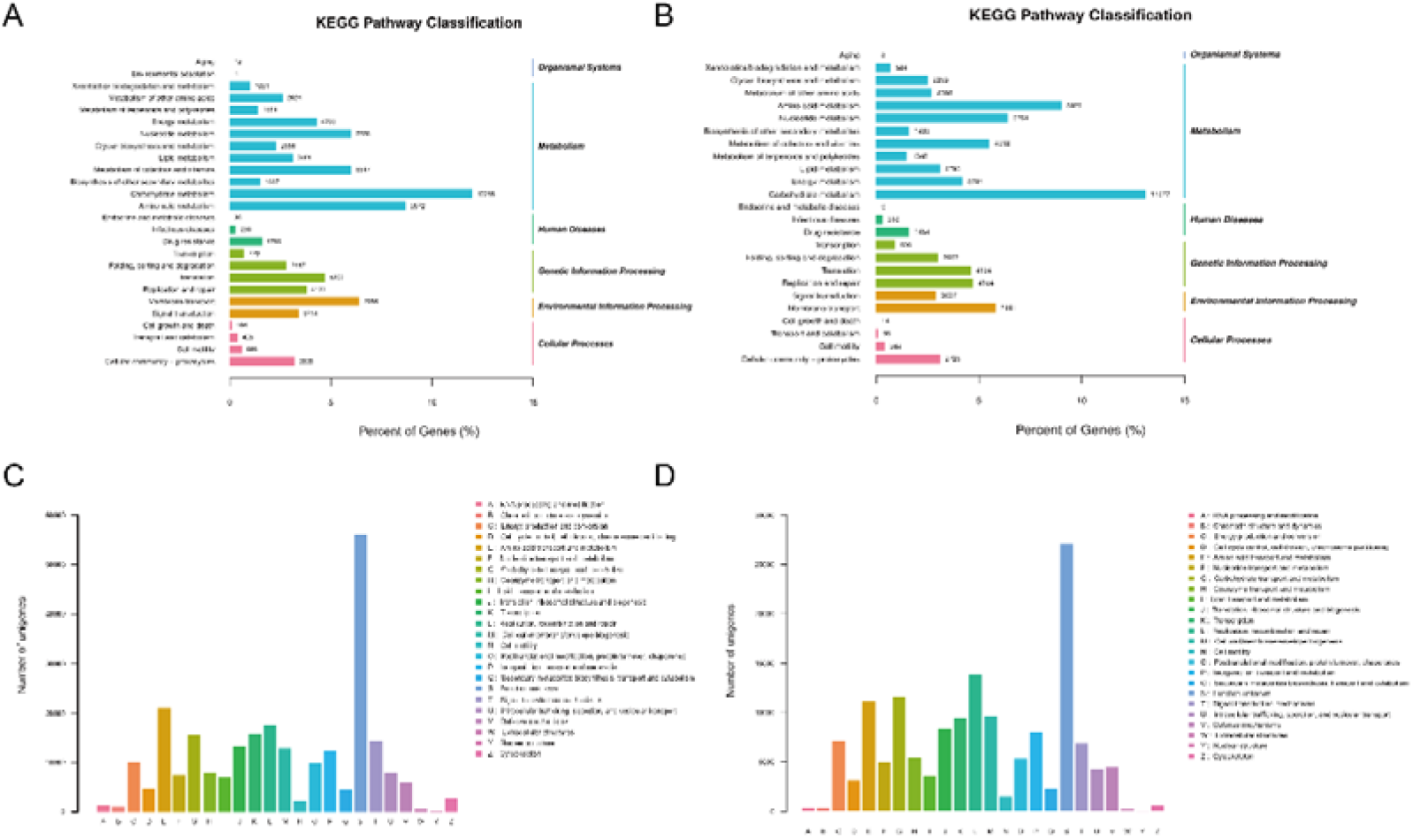
KEGG and evolutionary genealogy of genes (eggNOG) annotations of the intestinal metagenome and metatranscriptome in COVID-19 patients. (A, B) KEGG pathway annotations of the fecal metagenome and metatranscriptome, respectively. (C, D) eggNOG functional annotations for the gut metagenome and metatranscriptome, respectively.

### 2.6 Correlations between clinical indicators and fecal bacteria in COVID-19 group

Spearman correlation analysis was used to determine the relationship between the clinical data and bacterial microbiota (Fig. 8A). The result from MG shotgun data indicates seventeen bacteria were positively correlated with coagulation parameter APTT (Fig. 10A). Among them, *Prevotella* sp. BV3P1 is related with infectious parameter PCT. From MT data, *Firmicutes* CAG:103 and *Clostridiales* are positively correlated with PCT and platelets (Fig. 8B). In addition to basophils and lymphocytes, *Dorea longicatena* presents a negative correlation with other immune-related cells, such as neutrophils, leukocytes, and eosinophils. *Clostridia bacterium*, and *Eubacterium eligens* are negatively related to both basophil and lymphocyte.

**Figure 8.**
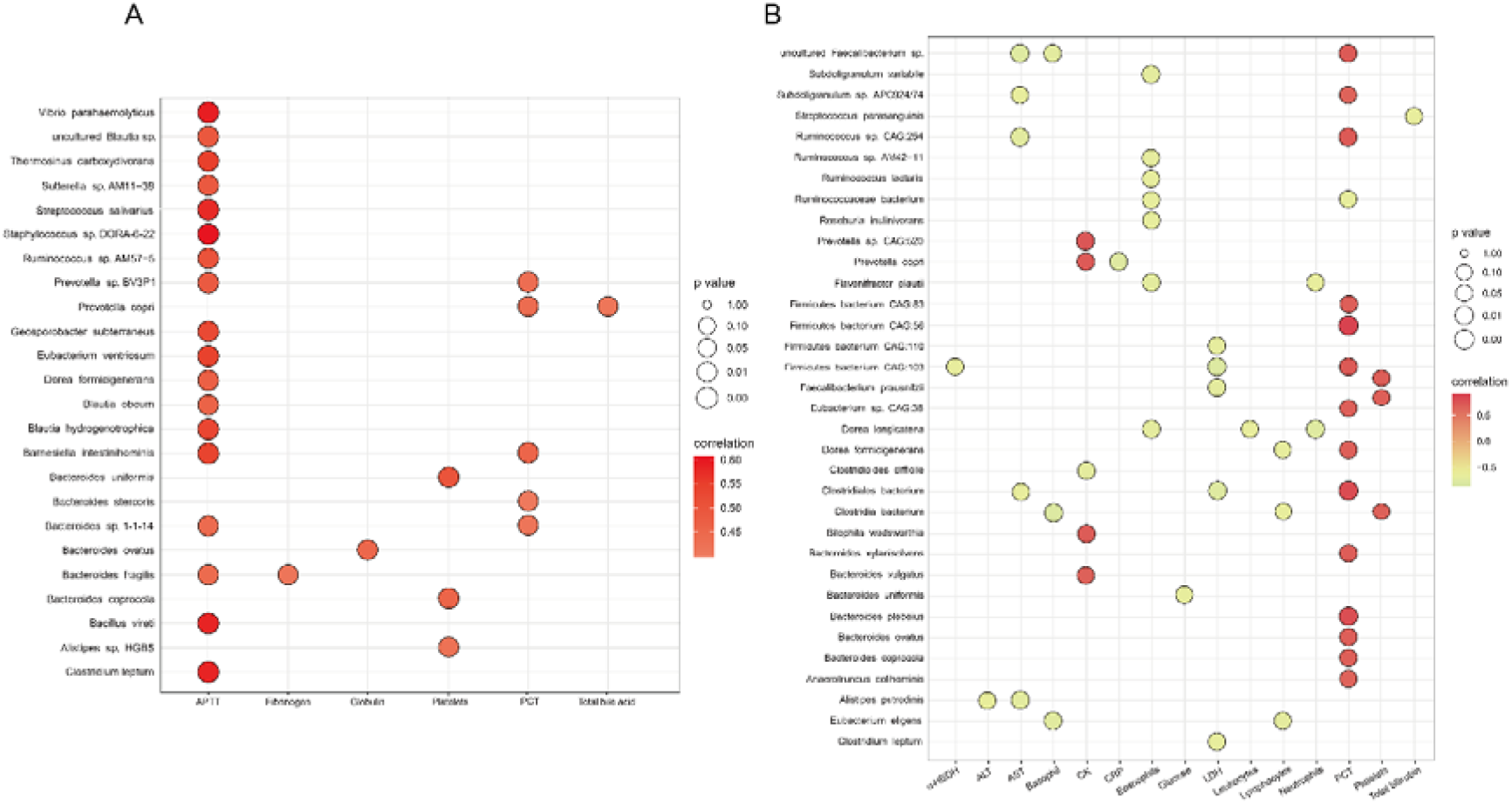
Relationship between clinical information of COVID-19 and species composition (abundance >0.1%) based on metagenomic (A) and metatranscriptomic (B) data, evaluated by spearman rank test. Only the results with P< 0.05 and absolute correlation coefficient over 0.4 are presented in the bubble plot. The degree of correlation is indicated by a color gradient from red (positive correlation) to green (negative correlation).

## 3. DISCUSSION

The decreased lymphocytes reflect possible bacterial infection and immune dysregulation in COVID-19 patients exposed to SARS-CoV-2 virus [25]. Shorter APTT is related to elevated risk of hypercoagulability and thromboembolism [26]. The clot waveform of APTT also suggested that COVID-19 patients might have distinctive abnormal coagulopathy [27]. High-level total bilirubin is generally considered a marker of abnormal liver metabolism and hepatitis, and there appears to be a significant relationship between hemoglobin levels and COVID-19 disease severity [28]. According to the mentioned results, the recruited COVID-19 patients might have liver damage, pathogen infection and blood system disorders. Disturbed gut microbiota may exacerbate COVID-19 severity through cytokine storm [29]. Thus, whether SARS-CoV-2 infection can interrupt the gut microorganism and affect COVID-19 progression should be confirmed. In this study, the following four main observations were performed to investigate the alteration of the gut microbial community under SARS-CoV-2 invasion.

The structure variation in the gut microbiota of participants was first examined. Our results revealed that there were significant differences in gut microbiota between COVID-19 patients and controls. The diversity and richness of the COVID-19 microbiome declined over time, which was considered related to respiratory viral infection [30]. Compared with health group, COVID-19 patients showed a greater diversity of microbiota between inter-individuals. Importantly, only a subset of patients returned to normal after discharge [18, 31, 32]. Subsequently, taxa abundance alternations were correlated to their properties. The F/B ratio was abnormally elevated in the COVID-19 group, suggesting a disease-specific shift in the overall microbiota composition of patients [33]. SARS-CoV-2 infection also resulted in the depletion of commensals in gut microbiota. Consistent with the previous study, the abundance of *Bacteroides* decreased and positively correlated with disease severity [34–36]. Among them ,*Bacteroides thetaiotaomicron* could down-regulate ACE2 expression level [37]. Moreover, it was metabolically complementary to butyrate-producing bacterium *Faecalibacterium prausnitzii* [38]. They together modulated the intestinal mucus barrier and could reduce SARS-CoV-2 virus load [18] [18, 34]. On the other hand, *Bacteroides fragilis* was involved in antiviral defense by inducing colonic plasmacytoid dendritic cells (PDC) [37]; whereas, *Bacteroides caccae* could regulate gut IgA levels [39]. In addition, *Bacteroides* could digest dietary and polysaccharides as host energy sources to promote the immune system [40]. A decrease in the abundance of indole-producing bacterium *Alistipes putredini* was also observed [41]. Indole is a precursor of tryptophan, and adequate tryptophan absorption is essential for intestinal antimicrobial peptides and α-defensins expression, thus maintaining epithelial barrier and endolymphocyte homeostasis [42]. *Alistipes putredini* was inversely associated with disease severity [43], possibly because it compensates for impaired tryptophan uptake due to ACE2 deficiency. The abundance of butyrate-producing bacteria *Lachnospiraceae*, *Blautia* and *Clostridium leptum* also significantly reduced [18]. It had shown that the decline of the butyrate producer is not conducive to COVID-19 recovery [18, 44, 45]. As one of the most important energy metabolism substrates for intestinal flora, butyrate plays a positive role in maintaining mucosal barrier, providing antiviral immune response and reducing inflammation [46] .

We also observed the accumulation of multiple opportunistic pathogens in the COVID-19 group. Increased abundance of *Actinomyce oris* aggravated COVID-19 severity [35]. Partial subspecies of *Escherichia coli* are important pathogens causing a variety of intestinal and parenteral infections [47]. *Streptococcus* was also found to be enriched in high SARS-COV-2 feature fecal samples [36, 43]. *Klebsiella pneumoniae* is common lung pathogen, and *Shigella* spp. are notorious gastroenteritis triggers [48, 49]. The enrichment of both *Eisenbergiella Tayi* and *Schaalia Odontolytica* increased the risk of bacteremia [50, 51], and the latter of which was discovered in various pulmonary infections and may be associated with the development of acute respiratory failure in patients [52, 53]. Further, some bacteria upregulated in the COVID-19 group were associated with inflammatory bowel disease, including *Subdoligranulum* spp., *Burkholderiales* sp. and *Gemmiger formicilis* [54–56]. Surprisingly, SARS-CoV-2 infection facilitated the growth of *Akkermansia muciniphila* and *Lactobacillus rhamnosus*. *Lactobacillus rhamnosus* is depicted as a potential co-infection microorganism along with SARS-Cov-2 [57]. *Lactobacillus* had been reported to aggravate mucosal inflammation, and interestingly, fecal samples from COVID-19 patients were rich in lactic acid [58]. *Akkermansia muciniphila* can improve intestinal barrier and provide host immune responses [59]. But, it had also shown that its abundance is positively correlated with H7N9 infection and disease severity [60]. This might due to the increased levels of Muc2, an essential ingredient for the growth of this bacterium, caused by respiratory virus infection [61]. Additionally, oral administration of this bacterium still inhibited H7N9 proliferation and improved clinical symptoms [62]. Thus, the endogenous increase of *Akkermansia muciniphil* in COVID-19 may be harmless.

Enrichment of pathogens relative pathways (i.e., *Staphylococcus aureus* infection, *Salmonella* infection, Pertussis and Bacterial invasion of epithelial cells) indicated the human gut is the site of extrapulmonary bacterial infection. The formation of *Escherichia coli* biofilms and their highly active expression support their association with COVID-19 severity. Galactose utilization could result in hypervirulent phenotype of *streptococcus pneumoniae* [63]. Dysregulation of the ABC transporter pathway implies that patients might be under toxic stress after exposure to SARS-CoV-2 [64].Besides, the accessory genomes of considerable pathogenic bacteria cover ABC transporters that contribute to antimicrobial resistance by multidrug efflux [65], further explaining antibiotic resistance pathway enrichment. Sphingolipids are progressively recognized as critical mediators in participation with inflammatory responses and multiple pulmonary diseases [66]. The biosynthesis of polyketide sugar unit and lipopolysaccharide could be related to oxidative stress state and risk of microbial translocation to systemic inflammation, respectively [67]. The results of gene annotation also indicated abnormal energy metabolism of intestinal flora in the COVID-19 group (Figure S12). The pathway of oxidoreductase activity hinted that active gut microbiota promoted energy-yielding biochemical reactions [68]. Three pathways of protein binding, single-stranded RNA binding and structural constituent of ribosome showed close relationships to metabolic processes, which demonstrated enhanced protein synthesis of gut flora as an extrapulmonary disease progression that cannot be underestimated [69]. Notably, impaired butyrate synthesis may indicate nutrient deficiencies in host cells. The increased degradation of neutral amino acids may be related to the consumption of ACE2, because the ACE2 level is closely related to the expression of the amino acid transporter B0AT1 [70]. Furthermore, branched-chain fatty acids derived from their degradation were related to obesity, metabolic syndrome and diabetes [71]. Taken together, intestinal microbiota impacted COVID-19 virulence accompanied with SARS-CoV-2 affected the intestinal microbiome, further participating in the pathophysiology of the host.

Lastly, as indicated from the MT data, the COVID-19 patients had various metabolically active microbiota. Among them, phylum *Verrucomicrobia* exhibited low transcriptional activity, which may be attributed to the low active state of *Akkermansia muciniphila*. The anti-inflammatory effect of *Akkermansia muciniphila* depends on the outer membrane protein Amuc_1100 [72]. Such a result may partially explain why the increased abundance of *Akkermansia muciniphila* did not endogenously alleviate COVID-19 and H7N9 progressions [62]. *Clostridium leptum* was reduced in the COVID-19 group, and its low metabolic activity further impeded its positive effect on disease progression. The active metabolism of *Prepotella copri* might also adversely affect COVID-19 development. In the upper respiratory tract, *Prevotella* was found to be positively associated with SARS-CoV-2 viral load [73]. As mentioned earlier, in MT data, the highly active ‘Cell wall/membrane/envelope biogenesis’ pathways participated in bacterial biofilm formation under SARS-CoV-2 infection, especially in *E*. *coli*. Moreover, active ‘membrane transport’ might be associated with antimicrobial resistance excretion.

Intestinal ACE2 is imperative in amino acid transport, tryptophan uptake, and antimicrobial peptide secretion [7]. The infection of SARS-CoV-2 not only consumes a lot of energy in the host cell, but also induces inflammation and reduces ACE2 expression. Correspondently, we observed that SARS-CoV-2 infection not only enhanced the metabolism of neutral amino acids in gut microbiota, leading to high oxidative stress and abnormal energy metabolism, but also contributed to enrichment of pathogens and the consumption of commensals. Notably, the malfunction of butyrate synthesis may suggest its adverse effect in disease progression [18, 34]. In turn, healthy gut microbiota can block the COVID-19 progression by providing host energy, maintaining mucosal barrier, and improving immune response, with *Bacteroides* likely a central player. Some limitations of the study should be mentioned. First, this is a single-center study with a moderate sample size, which does not apply to all COVID-19 patients. The corresponding relationship between SARS-CoV-2 infection and intestinal flora dysbiosis should be validated in a larger cohort, including subgroups at different stages of the disease. Though several species that may be central players for COVID-19 progression were discussed and reviewed (Table S9 and S10), meta-analysis from current multicenter and different studies to obtain universal conclusions is urgently warranted [18, 31, 32, 34, 35, 44, 58]. Moreover, this study depicted the alterations between patients at different stages of COVID-19, while no specific assessment has been conducted on the changes in COVID-19 microbiota and functions over time. Albeit we tried to control the variation degree between COVID-19 patients and the healthy controls, the alternations of gut microbiota may be influenced by other confounding factors, such as lifestyle, dietary habits, underlying diseases, complications, and clinical management. Lastly, the disease stage of COVID-19 at the time of stool sample collection is uncertain and there is a lack of information on pre-infection stool samples.

Collectively, this study further revealed alterations in the composition and function of active intestinal microflora in COVID-19 cases. Specific microbial biomarkers of COVID-19 patients were screened and correlated with clinical indicators. SARS-CoV-2 infection consistently exposes the intestinal microbiota to high oxidative stress and excessive inflammation state. Several key *Bacteroides* and butyrate-producing bacteria may play a pivotal role in mitigating COVID-19 progression. These results may deepen our understanding of how SARS-CoV-2 interferes with gut microbes and provide a treatment option for the fine-tuning gut microbiome in addition to the COVID-19 conventional treatment regimens.

## 4. METHODS

### 4.1 Epidemiological investigation

During the COVID-19 pandemic in China, all highly suspicious and confirmed cases in Lanzhou (36°03 ‘N, 103°40′ E) were admitted to hospitals abiding by the Infectious Disease Law. The First Hospital of Lanzhou University performed nasopharyngeal swabs-based real-time RT-PCR to screen 836 suspected patients to confirm SARS-CoV-2 infection. A total of 13 cases of COVID-19 and 24 cases of community-acquired pneumonia (CAP) were hospitalized. The patients diagnosed with SARS-CoV-2 infection by virology laboratory of this hospital were further confirmed by Lanzhou center for disease control and prevention (CDC) or Gansu provincial CDC. COVID-19 patients were classified as mild, moderate, or severe according to disease progression severity [18]. All pneumonia cases were recruited to this study for epidemiological investigation. 13 healthy subjects with age, BMI, and gender matching COVID-19 patients were also enrolled. Patients were cross-examined by hospital staffs pursuant to standardized questionnaires to generate clinical presentations and demographics. Medical records and laboratory results were reviewed to collect data on chest computerized tomography (CT), blood routine examination (WBC, NEUT, LYM, PLT, EOS, BAS HGB), coagulation function (APTT, d-dimer, PT, Fibrinogen), biochemical indicators (AST, ALT, TBA, TBIL, UA, CK, LDH, α-HBDH, AST) and inflammatory biomarkers (CRP , PCT). Examination of viral excretion from the nasopharyngeal and fecal samples was conducted by serial real-time RT–PCR.

### 4.2 Feces sampling and DNA/RNA extraction

The fresh fecal samples were collected by hospital staff using fecal collection tubes and a sterile stick, including 20 COVID-19 patient samples (13 baseline samples and 7 progression samples), 13 healthy samples, and 8 CAP samples (Fig. S1). All baseline samples of COVID-19 were collected during hospitalization (Fig. S2). Each fresh sample was delivered immediately from the ward to virology laboratory with ice packs where it was divided into aliquots of 1g and immediately stored at −80 °C until the next step. Next, DNA from all the above collected samples was extracted and RNA was extracted from 10 baseline fecal samples of COVID-19 group (Fig. S1). In brief, total bacterial DNA was extracted from the frozen aliquot of each fecal sample by using E.Z.N.A.® Stool DNA Kit (Omega, USA) following the manufacturer’s instruction. 1% agarose gel electrophoresis was employed to estimate DNA integrity. DNA purity was measured using Nanodrop spectrophotometer (Thermo Scientific, USA) and its concentration was determined using Qubit quantification system (Thermo Scientific, Wilmington, DE, USA). Total RNA was isolated and purified using E.Z.N.A.® Stool RNA Kit (R6828, Omega, USA) following the manufacturer’s procedure. The RNA amount and purity of each sample were quantified using NanoDrop ND-1000 (NanoDrop, Wilmington, DE, USA). The RNA integrity was assessed by Agilent 2100 with RIN number >7.0. The extracted DNA/RNA that conforms sequencing requirements was then stored at −80◻°C.

### 4.3 Metagenome and metatranscriptome sequencing and data analysis

DNA library was constructed by TruSeq Nano DNA LT Library Preparation Kit (FC-121-4001). In brief, DNA was fragmented by dsDNA Fragmentase (NEB, M0348S). The cDNA library was constructed by repairing the end of the DNA fragment, adding ‘A’ base to the blunt ends of each strand, adding sequencing adapters, fragments selection, and PCR amplification. For the extracted RNA samples, the Ribo-Zero™ rRNA Removal Kit (Illumina, San Diego, USA) was adopted to deplete rRNA and other host RNA sequences from total RNA. Subsequently, the left RNAs were fragmented and reverse-transcribed into cDNA. The cDNA library for sequencing was constructed as described in the above description. Finally, all cDNA libraries were sequenced on Illumina Novaseq 6000 (LC Bio, China).

The sequencing mode was performed with 150 bp paired end. Sequencing adapters and low-quality reads were filtered and trimmed from raw sequencing data by using cutadapt v1.9 and fqtrim v0.94 (sliding-window algorithm), respectively. Next, qualified reads were aligned to the human genome by employing Bowtie2 v2.2.0 to remove host contamination, followed by de novo assembly to construct the contigs for each sample by respectively applying IDBA-UD v1.1.1 and Trinity v2.2.0. All coding regions of contigs were predicted by using MetaGeneMark v3.26. And then, the contigs were clustered to by CD-HIT v4.6.1 to obtain unigenes. TPM was used to estimate the unigene abundance of a certain sample according to the aligned reads number of Bowtie2 V2.2.0. Then, unigenes were aligned against the NCBI NR database to obtain the lowest common ancestor taxonomy of them with DIAMOND v 0.9.14. Likewise, the GO/KEGG/COG annotations of unigenes were obtained.

### 4.4 Statistical analyses

The characteristics of the patients were described through demographics, epidemiological data, clinical signs and symptoms on admission, chest radiographic findings, laboratory results, treatment, and clinical outcomes of 2019-nCOV. Alpha diversity was calculated using QIIME v1.8.0. The principal coordinate analysis (PCoA) was used to assess beta diversity. The similarity analysis (ANOSIM) was conducted to compare the difference of microbiota structure between groups. The nonparametric MANOVA was performed to compare the effect size of subject metadata on microbiota composition. Differential bacterial taxa between groups were identified conducting Linear discrimination analysis (LDA) effect size (LEfSe) analysis, and taxon with an LDA◻>◻3.0-fold were considered significantly different. Using ANOVA with turkey multiple test correction to evaluate baseline microbiome related to COVID-19 severity (P <0.01 considered significant and F value acts as effect size). Spearman rank test was employed the correlation between clinical indexes and COVID-19 bacteriome. Wilcoxon test and Kruskal-Wallis test were used to analyze the differences of taxon/function pathways abundance.

### 4.5 Ethics approval

All participants provided written informed consents prior to starting the study. Research protocols were approved and supervised (LDYYL-2020-24) by the Institutional Review Board of the First Hospital of Lanzhou University and conformed to the ethical guidelines of the 1975 Declaration of Helsinki.

## ACKNOWLEDGEMENTS

This work was supported by Gansu Province COVID-19 (NCP) Science and Technology Major Project (2020) (No:20YF2FA008), Central University Basic Research Fund of China (No: lzujbky-2020-kb22), Science and Technology Planning Project of Lanzhou Chengguan District (No:2020JSCX0019), and National Natural Science Foundation Grants (No: 32070117 and 31870082). The author would like to thank all the medical staff working in the isolation ward of The Lanzhou University First Hospital. We also appreciate the assistance of Core Facility of School of Life Sciences, Lanzhou University.

## AUTHOR CONTRIBUTIONS

Tuoyu Zhou: Conceptualization; Writing - original draft; Validation and Formal analysis. Jingyuan Wu: Writing - original draft. YuFei Zeng: Genomic data analysis. Junfeng Li: Stool samples collection. Jun Yan : Epidemiological investigation. Wenbo Meng: Organization. Hawen Han: Revision. Pengya Feng: Methodology. Shuai Zhao: fecal DNA and RNA extraction. Ping Zhou: fecal DNA and RNA etraction. Yanling Yang: fecal DNA and RNA extraction. Ying Wu : Sample collection. Rong Han : Epidemiological data curation. Weilin Jin: Revision. Xiangkai Li, Yunfeng Yang and Xun Li: Supervision and Funding acquisition.

## COPETING INTERESTES

All authors declare that they have no known competing financial interests or personal relationships that could have appeared to influence the work reported in this paper.

## ABBREVIATIONS

COVID-19: Coronavirus disease 2019
SARS-CoV-2: severe acute respiratory syndrome coronavirus-2
APTT: Activated partial thromboplastin time
PT: Prothrombin time
AST: Aspartate aminotransferase
ALT: Alanine aminotransferase
CK: Creatine kinase
LDH: Lactate dehydrogenase
α-HBDH: α-hydroxybutyric dehydrogenase
CRP: C-reactive protein
PCT: Procalcitonin.

## Notes

### Competing Interest Statement

The authors have declared no competing interest.

